# GeneWhisperer: Enhancing manual genome annotation with large language models

**DOI:** 10.1101/2025.03.30.646211

**Authors:** Xiaomei Li, Alex Whan, Meredith McNeil, Samuel C. Andrew, Xiang Dai, Madeline Fechner, Cécile Paris, Rad Suchecki

## Abstract

Genome annotation is critical for understanding functional elements within genomes. Manual curation is a common practice for identifying the functions of genes, particularly those missed by automated annotation pipelines. However, this process is notoriously labour-intensive and time-consuming. In response to these challenges, we present GeneWhisperer, an innovative assistant system designed to facilitate the manual gene functional annotation process. Utilizing a large language model (LLM) agent, GeneWhisperer provides users access to tools appropriate for specific curation tasks in genome annotation. Featuring a conversational interface, GeneWhisperer fosters effective collaboration between human experts and AI. By synergizing the capabilities of AI with human expertise, GeneWhisperer offers a promising approach to advancing our understanding of gene functions and streamlining the gene annotation process.

## 1 Introduction

Manual annotation, or curation, of gene functions is a cornerstone in molecular biology and genomics. This detailed process requires systematic examination and interpretation of experimental data and scientific literature to determine the roles of individual genes within a genome. Thanks to advances in sequencing technology and assembly algorithms, the quality and completeness of genome assemblies have consistently improved. In this era of rapidly growing availability of genomic data, manual annotation remains crucial. It bridges the gap between raw sequences and functional insights, thereby deepening our understanding of the underlying genetic factors that contribute to biological phenomena.

Manual gene functional annotation is widely recognized as a time and labour consuming task. To address this challenge, a range of software tools have been developed to support the annotation process. These tools fall into several categories, each tailored to a specific aspect of manual annotation. For instance, Apollo (Dunn et al. 2019) and GeneDB (Manske et al. 2019) offer an online collaborative platform for gathering annotations from distributed experts. PubTator (Wei et al. 2019) and Textpresso (Müller et al. 2018) apply Natural Language Processing (NLP) techniques to automatically extract biomedical information from large volumes of literature, providing additional information for annotation. Finally, domain-specific tools such as BLAST (Altschul et al. 1990) and InterProScan (Blum et al. 2021) are instrumental in identifying sequence similarities and functional domains, respectively. Both are crucial for inferring the functions of genes.

However, current annotation practices often require the use of multiple software tools, which are not streamlined, thereby limiting their utility in supporting manual curation. Consequently, significant expert input and manual intervention are still required to curate and validate the information generated by these systems.

GeneWhisperer complements these existing tools by addressing key gaps in current annotation workflows. That is, it facilitates access to a range of annotation tools and enables the interaction between AI agents and human experts. An AI agent, also known as an autonomous agent, is an intelligent entity that perceives its environment, makes a plan, and takes actions to achieve its goals (Xi et al. 2023). AI agents could assist humans in various tasks and augment human intelligence (Richards et al. 2009; Peng 2021; Paris and Reeson 2024a). In recent years, the emergence of large language models (LLMs) has revolutionized the field of AI agents (Xi et al. 2023; Wang et al. 2024a). LLM-driven AI agents have shown significant potential to be applied to scientific discovery in multiple domains (Wang et al. 2024a; Boiko et al. 2023b; Szymanski et al. 2023). For example, ChemCrow (Bran et al. 2024) employs an LLM-driven AI agent to accomplish tasks across organic synthesis, drug discovery, and materials design. Finally, collaboration between humans and AI can leverage the strengths of both in practical settings (Paris and Reeson 2024b).

Our contributions in this work are twofold. Firstly, to the best of our knowledge, GeneWhisperer is the first LLM-driven AI system in the domain of manual genome annotation. This system offers the flexibility to integrate a variety of tools preferred by domain experts. Secondly, we propose an innovative integration of human and artificial intelligence within GeneWhisperer through a conversational system coupled with a semi-autonomous agent design. The semi-autonomous agent operates largely independently in task resolution, yet it retains the capability to seek assistance from human experts when necessary. We will detail the functionalities of GeneWhisperer in the subsequent section and elaborate on its use cases. Ultimately, we aim to make GeneWhisperer useful and trustworthy in biological research tasks, freeing up domain experts to focus on higher-level strategic thinking and creative problem-solving.

## 2 Related work

The process of manual genome annotation usually requires the use of multiple software tools that assist researchers (or curators) in analyzing and synthesizing data from diverse sources for informed decision-making. For instance, Gene Ontology (GO) annotation might involve using BLAST to identify homologous proteins, combined with NLP tools that retrieve relevant scientific literature. These tools help pinpoint mentions of proteins, their functions, and associated experimental evidence within the retrieved papers (Balakrishnan et al. 2013; Drabkin et al. 2012).

Traditionally, these tools were designed to perform specific tasks, each relying on distinct models tailored to their individual functions. However, the lack of integration between these tools results in inefficiencies, forcing annotators to frequently switch contexts and manually transfer data between systems.

In contrast, advanced LLMs can perform multiple tasks within a unified framework, ranging from text generation and translation to more specialized functions such as entity recognition and question answering (Supplementary Note 1). Furthermore, their capabilities extend to interacting with external tools and databases, enabling them to retrieve relevant information and seamlessly incorporate it into their outputs (Schick et al. 2023; Lewis et al. 2020). LLMs have shown remarkable potential to accelerate scientific discovery across a wide array of domains (Boiko et al. 2023a).

LLMs are increasingly being leveraged in the biological domain to tackle diverse challenges. For example, GeneGPT (Jin et al. 2024) utilizes an LLM to access data from NCBI web services and answer genomic questions. Similarly, DRAGON-AI (Toro et al. 2023) employs an LLM to generate new ontology components, while GeneAgent (Wang et al. 2024b) applies an AI agent to perform gene set enrichment analysis. Additionally, ChatGSE (Lobentanzer et al. 2023) harnesses an LLM to enhance biomedical analyses, including tasks like cell type annotation.

While these applications address certain biological challenges, they are not specifically designed to support manual curation tasks. Furthermore, they highlight the need for customizing LLMs to suit specific workflows. To address this gap, we have developed GeneWhisperer specifically tailored for manual curation, with a focus on GO annotation.

GO annotations describe the molecular functions, biological processes, and cellular components associated with gene products, providing a framework for understanding biological pathways and networks. These annotations, stored in the GO database (Gene Ontology Consortium 2021), are usually curated manually by experts. The curation process requires synthesizing information from multiple sources to support each annotation. GeneWhisperer enhances this process by providing annotators with synthesized, contextually relevant information from extensive biological databases and literature, thereby potentially reducing the manual effort required in data retrieval and integration tasks.

## 3 Methods

### 3.1 Overview

GeneWhisperer is a novel system that integrates an LLM-driven AI agent with domain-specific tools and facilitates interaction with human experts for gene function curation. Figure 1 illustrates the architecture of GeneWhisperer, which consists of an AI agent, a set of tools and a vector database. The AI agent selects tools relevant to user’s specific queries and interacts with the user. Through interactive dialogue, GeneWhisperer (1) identifies the most appropriate tool for the user’s query, (2) formulates the proper input for that tool, and (3) integrates the tool’s outputs in responses to the user. The tools perform essential tasks aiding manual curation, which we categorize into three main groups: searching scientific literature, extracting information from documents, and exploring public databases.

**Fig. 1.**
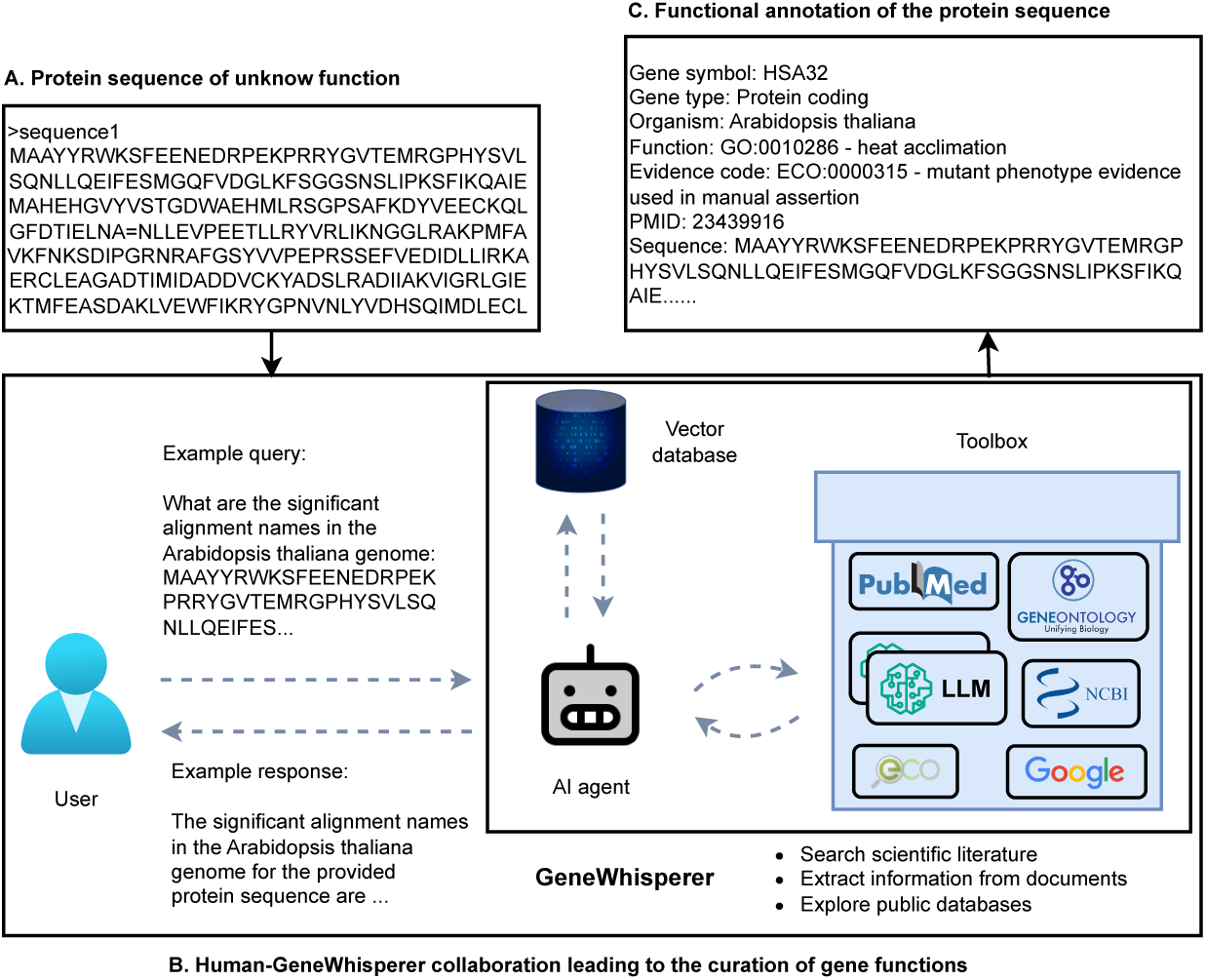
An overview of the manual curation process facilitated by GeneWhisperer. Rounded rectangles represent wrappers for tools such as NCBI BLAST, Google, an LLM, and others. Wrappers with an LLM icon represent an LLM configured with a customized prompt for information extraction tasks (Section 3.2.2). Other wrappers involve using an LLM with a customized prompt to facilitate access to data from external tools (e.g., BLAST), as detailed in Sections 3.2.1 and 3.2.3. The AI agent is introduced in Section 3.2.4.

These three categories encompass the essential manual curation tasks discussed in the literature (Balakrishnan et al. 2013; Hosmani et al. 2019; Ramsey et al. 2021). Accordingly, we have organized GeneWhisperer’s features into three primary functionalities. Each functionality may involve one or more tools, with each tool being independently wrapped and integrated into the toolbox. We have developed several LLM wrappers to handle different tasks (for more details, see Section 3.2).

Figure 1 illustrates how GeneWhisperer assists a user in manual gene functional annotation. In this scenario, a user identifies a protein of interest (Figure 1A) which has not been assigned a function by automated annotation pipelines. The user asks GeneWhisperer to align the protein sequence to a model species to identify homologous proteins. GeneWhisperer uses NCBI BLAST ^1^ to perform the sequence alignment task and provides the answer to this query (Figure 1B). To discover the function of this protein, the user may continually request GeneWhisperer to perform genome annotation tasks (more details in Section 3). Such human-AI collaborative effort results in the functional annotation of the protein sequence, as shown in Figure 1C.

In the following section, we will present each of these functionalities. We then introduce the AI agent of GeneWhisperer and its interactions with users in Section 3.2.3.

### 3.2 Main functionalities

#### 3.2.1 Searching scientific literature

Traditionally, researchers and curators manually navigate repositories such as PubMed (Williamson and Minter 2019) to identify scientific articles relevant to their work. Researchers first prepare specific keywords for PubMed searches. Once the PubMed search engine retrieves the relevant papers, they read through these documents (or abstracts) to determine the relevance to the research topics or questions at hand.

In contrast, GeneWhisperer streamlines this process by requiring users to input only their research question. Based on this question and previous chat history, an AI tool in GeneWhisperer automatically searches for relevant papers, summarizes information and provides citations to answer the user’s question. The chat history includes the user’s queries and the chatbot’s corresponding responses within the ongoing conversation. This interaction benefits users by allowing them to refine their questions iteratively and efficiently locate the desired literature. If no chat history exists when a dialogue starts, the system relies solely on the current question.

We have integrated an LLM-powered PubMed wrapper into the toolbox to facilitate this functionality. This wrapper uses an LLM to analyze the user’s question, generate relevant keywords, and conduct PubMed searches (Figure S1). To enhance its performance, we utilize a prompt engineering technique, i.e., few-shot learning, to guide an LLM in performing specific tasks. Few-shot learning involves providing the model with a small number of example inputs and corresponding outputs within the prompt, enabling it to execute the desired task effectively (more details in Supplementary Note 2). The wrapper then provides an answer based on the retrieved literature (default setting: top 5 ranked papers published within the last 10 years). For more details, please refer to Supplementary Note 3.

#### 3.2.2 Extracting information from documents

Through the literature searching functionality in GeneWhisperer, users can refine their search results to identify papers of interest. For those selected papers, users may seek to gain deeper insights and detailed information. GeneWhisperer supports this process by offering a suite of tools to extract and summarize information from a document (Figure 2).

**Fig. 2.**
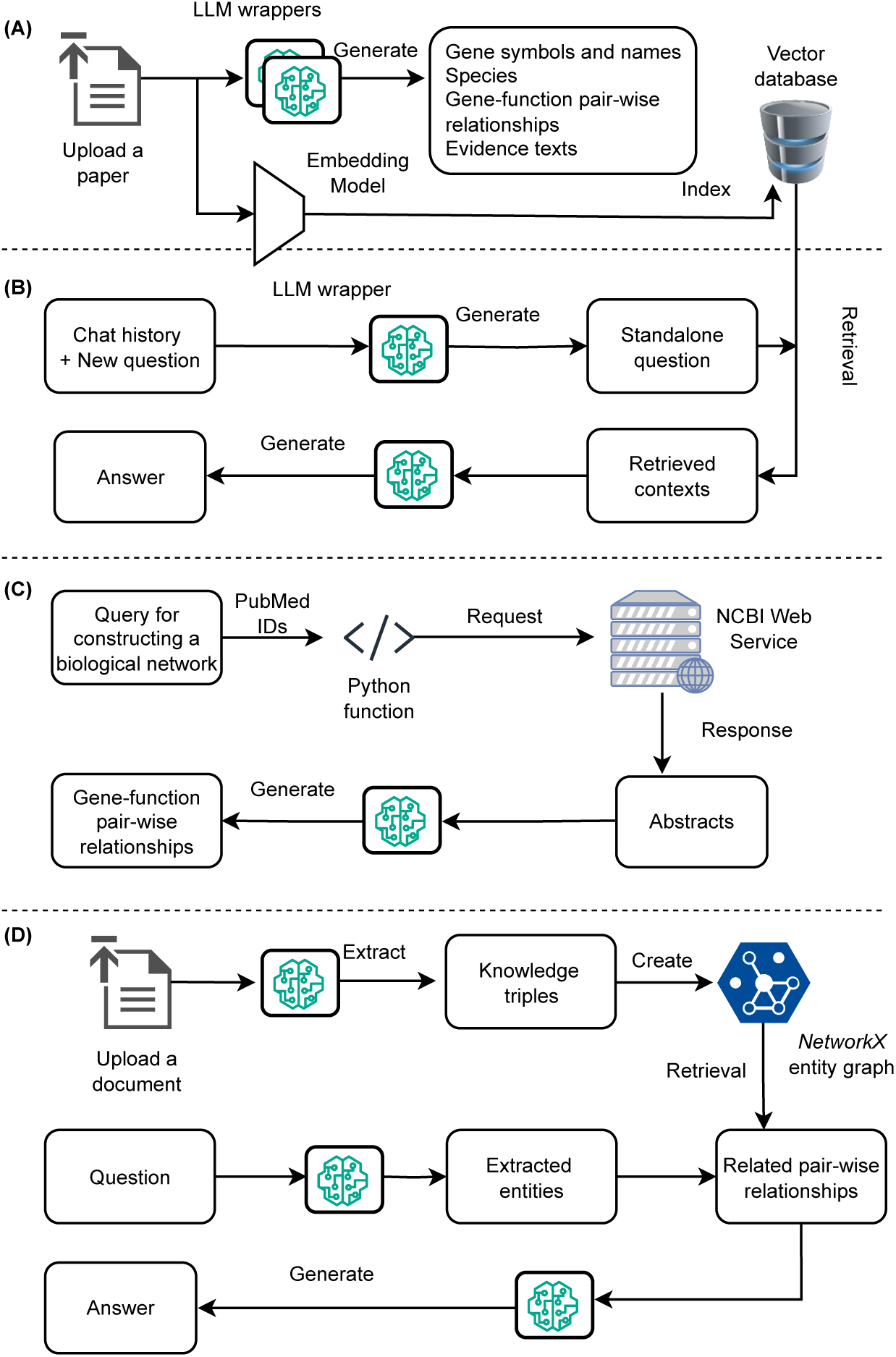
Tools for extracting and summarizing information from documents: (A) Retrieve metadata and index the paper into a vector database. (B) Answer user queries based on information within the paper. (C) Construct a biological network from a list of scientific papers. (D) Answer user queries using information derived from the biological network. Rounded rectangles represent either input or output text. Arrows illustrate the processes and actions performed by the AI system behind the scenes. The icons highlight key components such as the embedding model, vector database, LLM wrappers, NCBI web API service, and knowledge graph, demonstrating how information is processed for the user. An LLM wrapper refers to an LLM with a customized prompt, as detailed in Supplementary Notes 4-5.

Users can upload a paper of interest, GeneWhisperer then generates metadata about its abstract and indexes the full document in a vector database (Figure 2A). The vector representation is calculated by an embedding model (e.g., *OpenAIEmbeddings* in *LangChain* (Chase 2022)), which processes the document to extract semantic information and encode it as high-dimensional embeddings. These embeddings capture contextual relationships and allow for efficient similarity searches and downstream analyses. For tasks such as question answering, the embeddings serve as a bridge to identify and retrieve the most relevant portions of the document, which are subsequently interpreted by an LLM to generate precise responses.

The metadata consists of standardized summaries used for manual curation, including gene definitions, species, functional relationships among a wide range of biological entities, and supporting evidence. In this context, evidence refers to any relevant information that supports the conclusions in the scientific paper, which can arise from experimental (such as wet lab) techniques or computational (*in silico*) methods. All this information is synthesized using the LLM wrappers (an LLM with customized prompts, detailed in Supplementary Note 4).

The metadata provides initial gene annotations from the paper of interest. Users may need to dive further into the paper to verify the annotation or find more gene functions in other sections. Instead of manually reviewing the entire text of the lengthy paper, users can ask GeneWhisperer questions, and it will provide answers based on the information within the paper (Figure 2B). To enable this capability, we integrate an LLM into a Retrieval-Augmented Generation (RAG) framework (Gao et al. 2023). RAG enhances the Question-Answering (QA) process by combining a vector-based retrieval engine, which identifies the most relevant contexts from the document, with the generative capabilities of the LLM. This approach ensures that the responses are not only contextually accurate but also grounded in the original content of the paper, providing users with reliable and detailed answers tailored to their queries.

Annotating genes with functional relationships is an important part of manual curation, as shown in the Apollo system (Dunn et al. 2019). We also implemented a tool in GeneWhisperer to extract functional relationships from a list of research papers (Figure 2C). A functional relationship is represented as a triplet (*E*1*, E*2*, re*), where *E*1 and *E*2 are biological entities, and *re* is the relationship between them. GeneWhisperer first retrieves the abstracts associated with the provided PubMed Identifiers (PMIDs) and then applies a method outlined in the literature (Fo et al. 2023) to prompt an LLM for extracting gene-function pairwise relationships mentioned in the abstracts. Additional technical details of this process are available in Supplementary Note 5.

These functional relationships can be formatted as a knowledge graph, with biological entities as nodes and the relationships as edges mapping the interactions between entities within biological systems. The knowledge graph is stored locally as a file and indexed using a wrapped *NetworkX* entity graph instance (via the *GraphIndexCreator* function in *LangChain*), enabling further network analysis and visualization. The indexed knowledge graph can be utilized by GeneWhisperer to answer graph-related questions. Specifically, it extracts entities from a user’s query, retrieves relevant relationships from the indexed knowledge graph, and generates answers based on the retrieved information. As shown in Figure 2D, users also have the option to upload a pre-existing knowledge graph instead of generating one through the system, providing flexibility for integrating external data into their analyses.

#### 3.2.3 Exploring public databases

In manual curation, users often search gene-related information from National Center for Biotechnology Information (NCBI) databases (Sayers 2010), gene ontology terms from the Gene Ontology (GO) database (Gene Ontology Consortium 2021), and evidence terms from the Evidence and Conclusion Ontology (ECO) database (Nadendla et al. 2022). GeneWhisperer addresses these types of user requests by leveraging the AI agent to navigate these diverse databases.

We have developed LLM wrappers for each database to facilitate effective interaction by the AI agent. NCBI serves as a critical repository for biological data, encompassing various databases such as gene, SNP, OMIM, and sequence databases (e.g., non-redundant sequence database). To address queries related to gene aliases, gene SNPs, and gene-disease relationships, we employ methods outlined in GeneGPT (Jin et al. 2024) to prompt an LLM to interact with the NCBI gene, SNP, and OMIM databases, respectively. We prompt an LLM to generate the corresponding NCBI web API endpoint, which is a URL address serving as the point of contact between a user application and the NCBI API server (Sayers 2010). This API endpoint can be accessed using Python’s *requests* function to retrieve data (e.g., summary information about a gene). These data are appended to the LLM’s prompt, enabling it to generate the final answer to the user’s question. This process is demonstrated in examples where, through few-shot learning, an LLM generates web addresses for data crawlers and answers user questions based on crawled data. We employ a similar technique for questions related to GO or ECO, using an LLM wrapper to call the quickGO web APIs to extract information from these databases.

GeneGPT provides controlled outputs formatted according to predefined examples. However, its application to BLAST tasks can quickly exceed the token limits of LLMs. To address this, we employ RAG to wrap the BLAST tool, similar to the process shown in Figure 2C. First, an NCBI BLAST endpoint is generated for the user’s query sequence. The Python *request* function is then used to retrieve data from this endpoint. The BLAST result data are indexed in a vector database, and a retrieval engine extracts relevant information from this database. Finally, an LLM generates an answer based on the retrieved results.

Additionally, we can leverage other search engines to expand search capabilities in GeneWhisperer beyond NCBI, GO and ECO databases. For instance, GeneWhisperer may use a Google search wrapper (e.g., *SerpAPIWrapper* from *LangChain*) to identify the most relevant web pages in response to a user’s query.

In summary, we have built wrappers to access NCBI, GO, ECO, and Google resources that may be useful during manual curation. Each wrapper tool is an independent module, and this modular design facilitates the easy integration of new tools into the GeneWhisperer framework, ensuring both flexibility and adaptability.

The AI agent and its interactions with users We employ the *LangChain* framework to implement the AI agent (Supplementary Note 6). *LangChain* supports a wide variety of components including prompt templates, LLM models, document loaders, information retrieval tools, vector databases, and memory management. In GeneWhisperer, vector databases are used to index and store user-uploaded documents, providing domain knowledge bases for the AI agent. Memory management involves storing and retrieving past conversations to enhance the contextual awareness and engagement of an AI system. Through the *LangChain* framework, our AI agent is integrated with an array of tools and external resources, optimizing the output from a given input question.

Recent research indicates that existing autonomous AI agents experience performance degradation when tackling practical tasks (Shi et al. 2024; Yuan et al. 2024). However, incorporating human-AI interactions can significantly improve the effectiveness of AI agents in various applications (Xiao and Wang 2023; Yang et al. 2023). Accordingly, the AI agent in GeneWhisperer is designed as a semi-autonomous agent that seeks human participation in the reasoning process if needed, fostering collaboration between humans and AI.

To facilitate human-AI interactions, GeneWhisperer is designed with an intuitive conversational user interface (Figure 3). Users primarily communicate with GeneWhisperer in natural language. They can also click one of the three buttons on the interface depending on their requirements. The agent-based system setup enables a wide range of functionality with just a few buttons.

**Fig. 3.**
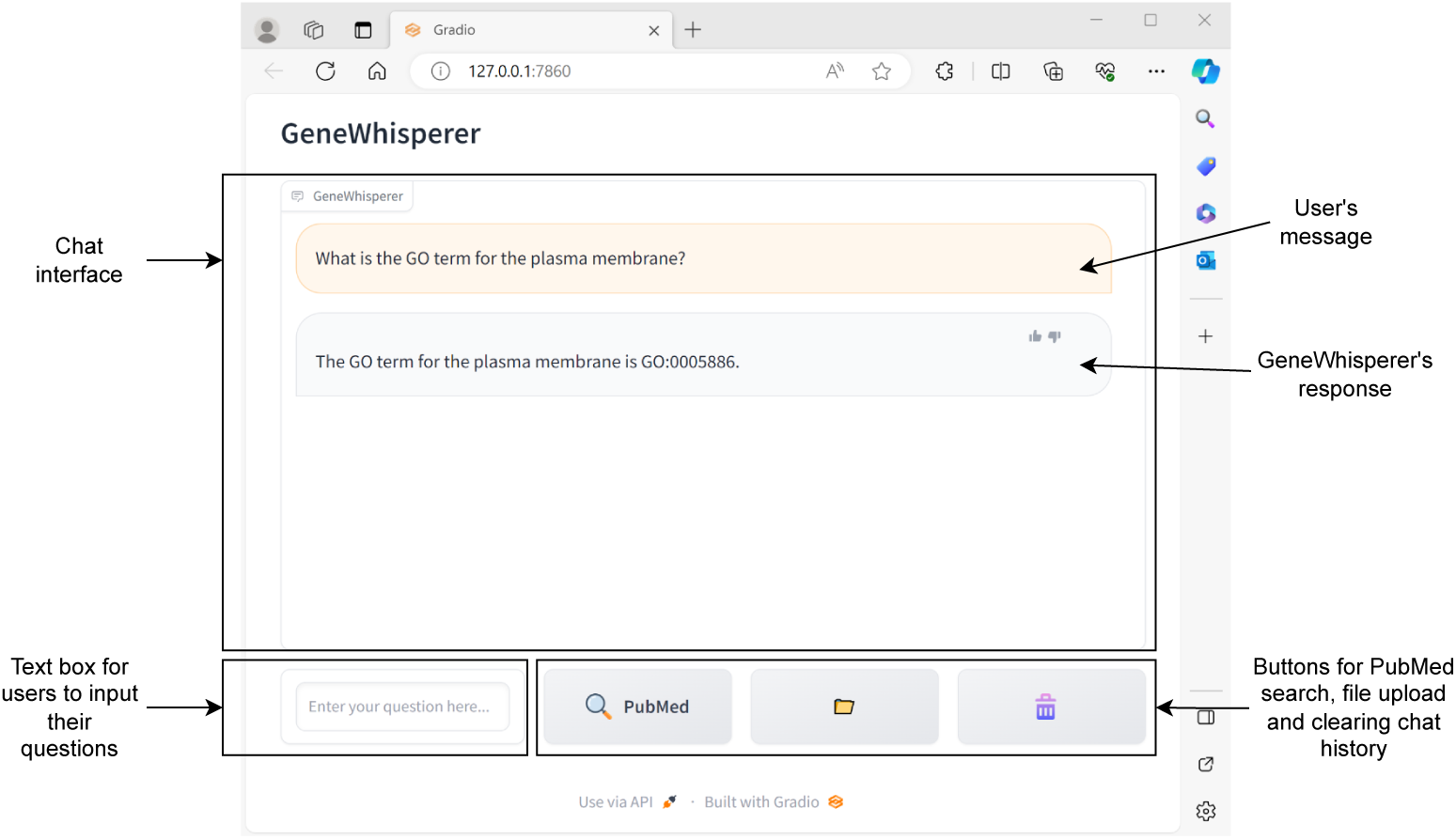
The user interface of GeneWhisperer. The ongoing conversations are displayed in the chat interface area, showing both the user’s messages and the GeneWhisperer’s responses. A text box allows users to enter their questions, triggering GeneWhisperer to execute the AI agent. Given the frequent use of PubMed by users, we have included a direct PubMed search button in the user interface for convenience. There are two other buttons, one for file upload (of e.g., a scientific paper or a file containing a knowledge graph), and one for clearing chat history.

We show how a user can use GeneWhisperer in Figure 4 and demonstrate how this human-AI collaboration improves the performance of the AI agent while supporting the human in a specific use case, as detailed in Supplementary Note 7. This example also illustrates the semi-autonomous nature of our agent, as it seeks human assistance in the reasoning process.

**Fig. 4.**
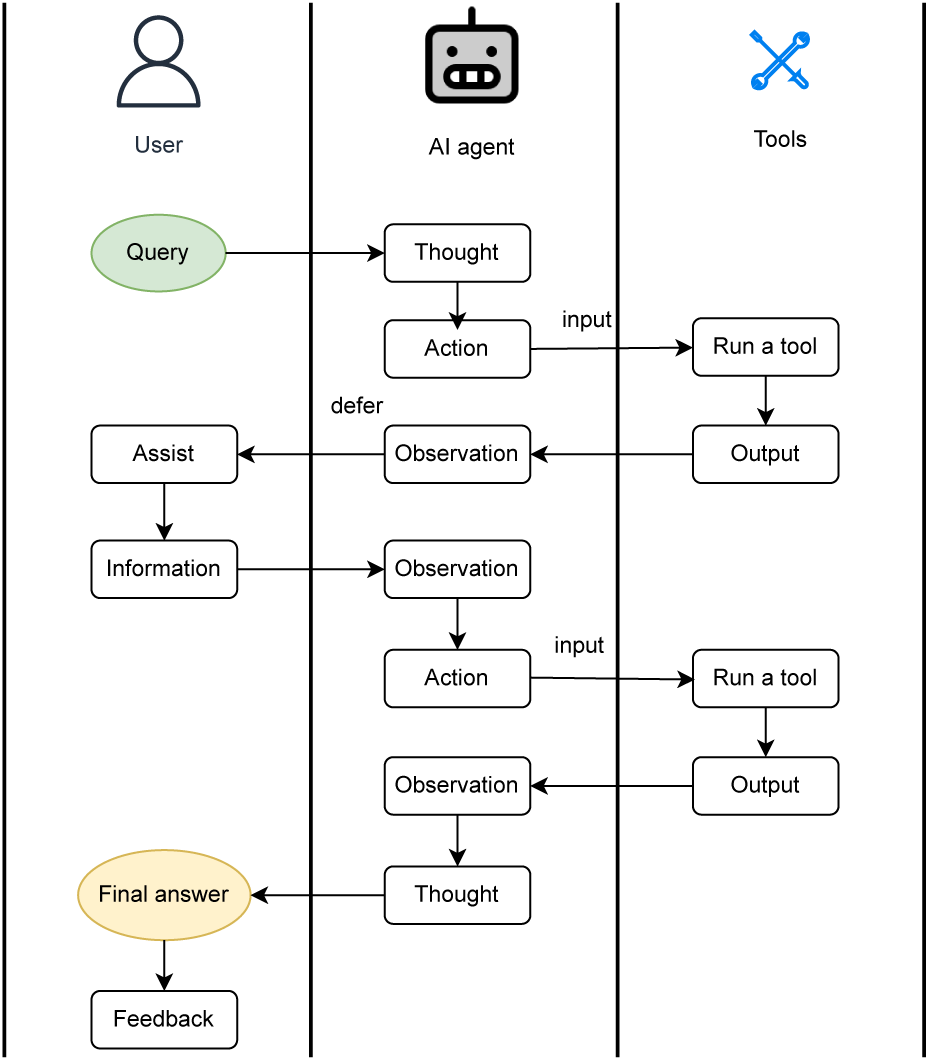
Example of a human-AI collaboration workflow using GeneWhisperer. ➀ a user poses a query; ➁ the AI agent processes the query and executes a tool; ➂ the AI agent requests assistance from the user; ➃ the user provides additional information; ➄ the AI agent analyzes the human input and chat history, then executes potentially another tool; ➅ the AI agent analyzes the tool outputs and chat history, and presents a final answer to the user. Processes ➃-➅ may be repeated several times to refine the response.

## 4 Results

### 4.1 Prompt engineering significantly improves NCBI searches

Prompt engineering is one of the strategies that help improve the quality of responses given by LLMs. Prompts are a form of programming that uses natural language to give context, instructions and rules which help to guide an LLM towards the desired output (White et al. 2023). The effectiveness of a prompt depends on a multitude of factors, from subtle alterations of syntax and semantics (Fu et al. 2022) to patterns modelled on human decision making (Yao et al. 2023).

We employed an iterative prompting method to enhance the accuracy of tasks involving the navigation of NCBI databases. Specifically, we started with the baseline prompts used in GeneGPT, scored the responses, evaluated the sources of error, identified patterns, designed alternative prompts based on the error analysis (Supplementary Note 8), and then tested them in subsequent trials.

The baseline GeneGPT prompts consist of three components: a contextual statement, two documentations, and four one-shot demonstrations (Jin et al. 2024). The contextual statement informs an LLM that it will be tasked with answering genomic questions and that it can utilize the NCBI web APIs to do so. The documentations provide general information about using NCBI web APIs and executing BLAST searches. The four demonstrations exemplify how to use each documentation in context by taking a question from sample genomic QA tasks and listing the steps needed to obtain the provided gold standard answer. The demonstrations are for 1) Gene alias, 2) Gene SNP association, 3) Gene disease association and 4) DNA sequence alignment to the human genome. The genomic question-answer pairs are derived from the GeneTuring (Hou and Ji 2023) and GeneHop (Jin et al. 2024) datasets.

We iteratively modified the demonstrations for NCBI web API endpoint generation and changed the documentation based on the error analysis. We conducted experiments in gene alias, gene location and SNP-associated gene function tasks which had low scoring in GeneGPT compared with other tasks. We found that the refined prompt significantly improved the LLM’s performance on these three genomic QA tasks, each consisting of 50 question-answer pairs. The accuracy indicating scores increased from 0.69, 0.66, and 0.02 to 0.97, 0.95, and 0.66 for the gene alias, gene location, and SNP-associated gene function tasks, respectively. Please refer to more experimental details in Supplementary Note 8 and Supplementary Data 1-3.

### 4.2 GeneWhisperer accurately identifies sequence alignments

Accurately identifying significant sequence alignments is a critical task in genome annotation, particularly when analyzing unknown sequences. To evaluate GeneWhisperer’s ability to perform this task, we adapted the DNA sequence alignment question-answer pairs from the GeneTuring dataset to suit our objective.

The original alignment task evaluates AI models on their ability to determine which chromosome a DNA sequence aligns to in the human genome. The question was created by appending the phrase *“Align the DNA sequence to the human genome:”* before a DNA sequence. The corresponding answer is a chromosome name.

For our evaluation, we reformulated the questions to fit our purpose by appending the phrase *“What are the significant alignment names in the human genome:”* before the DNA sequence. The significant alignment names obtained through BLAST searches served as the gold standard answers. Using the same DNA sequences from the GeneTuring dataset, we generated 50 question-answer pairs to test GeneWhisperer’s performance. Note that this evaluation dataset was created specifically for testing purposes, and GeneWhisperer is capable of analyzing sequences from other species as well.

We evaluated GeneWhisperer on three key aspects: whether it correctly called the appropriate BLAST API with the correct input DNA sequences on its first attempt, whether its outputs adhered to the RAG framework format (Supplementary Note 6), and whether its final answers were accurate. The GPT-3.5-turbo model was used as the backend LLM for GeneWhisperer during the evaluation. Our results showed that GeneWhisperer achieved 100% accuracy in calling the correct BLAST API with the appropriate input DNA sequences on its first attempt. However, for one question, GeneWhisperer failed to provide the correct final answer, resulting in a 98% accuracy rate. In this instance, although GeneWhisperer used the correct BLAST API and input DNA sequences, the alignments in the BLAST results did not correspond to the human genome. Consequently, GeneWhisperer attempted to identify human homologs by searching for one of the gene names from the BLAST results using Google. When this search failed, it sought human assistance. While this demonstrates the potential of AI agents to leverage multiple tools to solve complex problems, this specific use case was outside the scope of our testing and was not pursued further. In terms of output format, only 44 out of 50 responses followed the strict RAG framework format, yielding an 88% accuracy rate. However, deviations from the output format did not affect task completion accuracy in our evaluation. For additional experimental details, please refer to Supplementary Data 4.

### 4.3 GeneWhisperer leverages domain knowledge for accurate genome annotation

While the guidelines for manual curation may vary across projects, certain common steps can be followed. Drawing on the literature (Balakrishnan et al. 2013; Drabkin et al. 2012), we adapted the annotation pipeline used in the human gene function annotation project by the UCL Institute of Cardiovascular Science ^2^. In this pipeline, manual curation begins with a given gene sequence, alias, or symbol. Through a systematic querying process, human annotators gather and curate information for use in GO annotation. They then evaluate and justify this information before creating final annotations for the given gene (Figure 5).

**Fig. 5.**
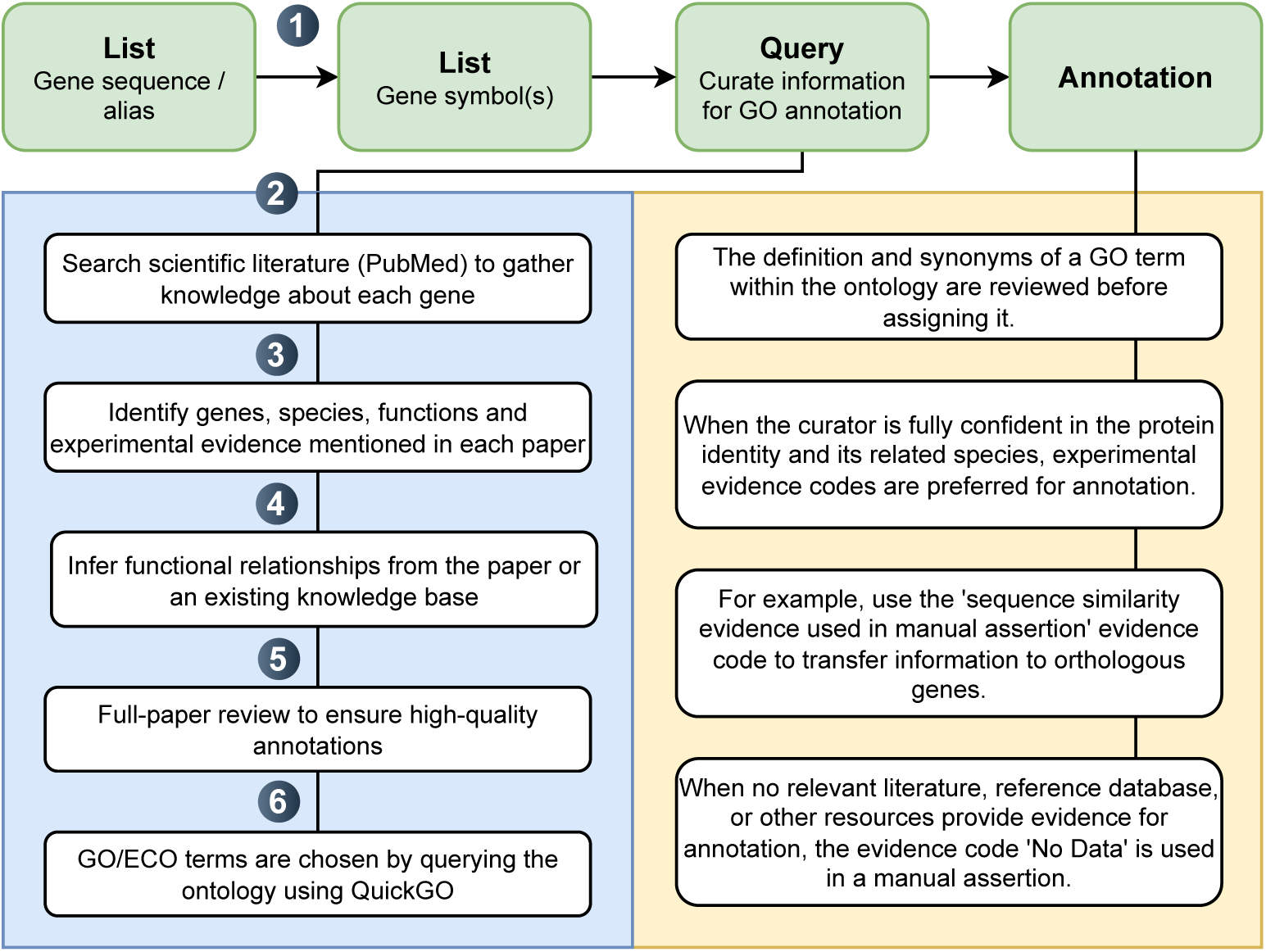
A pipeline for manual gene (or gene product) function annotation. GeneWhisperer assists with Steps 1-6, generating information for the user to review. The final annotations are then determined based on the user’s expertise and judgment, as represented by the processes in the unnumbered text boxes on the right.

We present examples of GeneWhisperer assisting in the manual curation process, specifically following Steps 1 to 6. The GPT-3.5-turbo model serves as the backend LLM for GeneWhisperer in generating these responses. For comparison, we also include the original responses generated directly by GPT-3.5-turbo.

**Step 1.** Search homologous proteins and gene symbols for a target protein

Genome annotations frequently include many uncharacterized proteins, especially in non-model species. Identifying the functions of these proteins often begins with searching for their homologs in closely related and presumably annotated species’ genomes. To achieve this, GeneWhisperer can be used to align the sequence of the query protein to a relevant database which includes appropriate model species data. The queries and responses are provided in Supplementary Data 5. The results indicate that GPT-3.5-turbo is unable to answer this question due to its lack of access to BLAST to address the problem. By leveraging its tool-use capabilities, GeneWhisperer successfully employs the BLAST search tool to complete this task.

To review these proteins in the literature and various databases, it is essential to identify the gene names and symbols coding for these proteins. Our results demonstrate that GPT-3.5-turbo provides an incorrect answer to this question. In contrast, GeneWhisperer successfully addresses it by utilizing NCBI web APIs to query the NCBI gene database. Subsequently, the most similar protein is selected for detailed further study.

**Step 2.** Search scientific literature to identify papers of interest

In the manual curation process, the functions of HSA32 are transferred to the target protein. To achieve this, it is essential to identify and validate the functions of HSA32 as documented in the scientific literature. We present two example queries that users might use to search for relevant papers. While GPT-3.5-turbo provides responses to these queries, their accuracy cannot be verified without references. In contrast, GeneWhisperer delivers answers accompanied by cited papers, enabling users to conduct a more in-depth analysis.

**Step 3.** Identify genes, species, functions and experimental evidence mentioned in each paper

GPT-3.5-turbo is not designed to answer user questions based on specific papers. In contrast, GeneWhisperer utilizes LLM wrappers to extract genes, species, functions, and experimental evidence mentioned in each paper (methods detailed in Section 3.2.2). These metadata provide users with preliminary information, helping them decide whether to conduct a deeper analysis of the paper.

**Step 4.** Infer functional relationships from the paper or an existing knowledge base

Once the user has the information about the putative function, they may want to explore the gene’s regulatory relationships. Such genes regulatory relationships, along with other functional relationships, can be organized into a biological knowledge graph. Relationships presented in such knowledge graphs can aid in inferring gene functions. For example, a gene’s function can be deduced through its direct physical interactions with other gene products or through associations between genes.

In this task, we use GeneWhisperer to construct a knowledge graph based on the abstracts of six papers related to HSA32. To accomplish this, we ask GeneWhisperer, *”Can you build a biological network based on PMIDs 23439916, 24548003, 25711624, 24520156, 30080608, and 25013950?”* GeneWhisperer extracts all the functional relationships within the abstract of these papers and indexes the knowledge triplets in a knowledge graph. In contrast, GPT-3.5-turbo responded that it could not construct a biological network using the provided PubMed IDs.

As mentioned earlier, the knowledge graph serves as contextual support for GeneWhisperer to answer questions related to functional relationships. Users can also upload their own knowledge graph file (e.g., interaction data from the IntAct molecular interaction database (Orchard et al. 2014)), which GeneWhisperer will then index. GeneWhisperer can infer its answers from the indexed knowledge graph, regardless of whether it was generated by GeneWhisperer or uploaded by users. In contrast, GPT-3.5-turbo answers such questions based on its pre-trained data rather than using domain-specific context.

In this use case, we demonstrate the usefulness of knowledge graphs in manual curation, highlighting the importance of integrating this functionality into our tool. In fact, question answering over knowledge graphs has recently emerged as a significant research area. Knowledge graphs are particularly effective for answering questions that require multiple steps of reasoning (Chakraborty 2024).

**Step 5.** Full-paper review to ensure high-quality annotations

Reviewing the full paper gives important context for users to generate high quality annotation. For instance, groups of biological entities may be mentioned in the abstract without detailed names. Consequently, users need to identify the specific names for these entities. Moreover, users may read more details about the experimental results to justify the annotation. Similar to the previous task, GeneWhisperer provides more accurate answers than GPT-3.5-turbo by relying on factual information from the paper.

**Step 6.** Gene annotations with controlled vocabularies in GO and ECO

Curators usually employ controlled vocabularies for gene annotations, ensuring consistency and accuracy in their work. This standardized approach facilitates the understanding and comparability of genetic data across different studies. GeneWhisperer effectively assists with this task by leveraging QuickGO, whereas GPT-3.5-turbo often produces incorrect answers based solely on its trained data.

## 5 Discussion

Manual annotation is time-consuming, as it involves curating gene annotations from diverse sources. To address this, we developed GeneWhisperer, a novel system that integrates an LLM-driven AI agent with domain-specific tools and interacts with human experts for gene function curation. GeneWhisperer has the potential to greatly benefit the biological research and curator communities. For instance, high-quality curated gene functions can provide biologists with valuable insights to shape their hypotheses and design experiments more effectively. These curated gene functions also serve as a golden standard for researchers developing computational tools to accurately predict gene functions (Zhou et al. 2019).

Our experimental results show that an LLM agent powered by domain-specific tools outperforms an standard LLM model (GPT-3.5-turbo in this study) in genome annotation tasks. These domain-specific tools are seamlessly integrated into a unified framework, allowing the AI agent to use them effectively. Additionally, we enhanced the LLM’s performance by incorporating task-specific instructions and examples through prompt engineering. The results demonstrate that prompt engineering significantly improves the effective utilization of these tools. Furthermore, embedding the LLM with domain knowledge (whether derived from tools such as BLAST or scientific literature) within a RAG system enables the AI agent to accurately answer user queries.

We have demonstrated the potential benefits of utilizing an AI agent and fostering human-AI collaboration in the manual curation process for gene function annotation. However, the manual curation process can be more complex in real-world applications. For instance, while BLAST is a powerful tool for sequence alignment, it has limitations in function prediction. The underlying assumption when using BLAST for function prediction is that proteins with similar sequences have similar functions, which is not always valid in biology contexts (Al-Fatlawi et al. 2023).

To address these challenges, additional tools can be integrated to verify whether two genes share similar functional annotations. Examples include AlphaFold3 (Abramson et al. 2024) for structural prediction, InterProScan (Blum et al. 2021) for identifying conserved domains and functional sites, and OrthoDB (Kuznetsov et al. 2023) for ortholog identification and validation. The combined outputs from these tools provide biocurators with valuable insights, helping them assess whether sequence similarity is a reliable basis for inferring the function of a gene product (Balakrishnan et al. 2013).

Challenges in manual curation remain, even with the assistance of GeneWhisperer. First, GeneWhisperer and similar AI assistants depend heavily on the performance of LLMs, both in general and within specific domains. For example, the reasoning and language understanding capabilities of LLMs are crucial for accurately interpreting user queries and selecting the appropriate tools to address them (Xi et al. 2023).

RAGs and prompts are two strategies for enhancing LLM performance. RAG engines retrieve relevant texts to provide context for the LLMs, thereby improving their accuracy in answering questions. The choice of RAG engines plays a significant role in determining LLM performance (Cuconasu et al. 2024). Similarly, well-crafted prompts with clear and specific instructions tailored to domain-specific tasks guide LLMs to produce more focused and accurate outputs. However, evaluating RAG systems and prompts presents unique challenges due to their reliance on dynamic knowledge sources. At present, we rely on human evaluation due to the absence of benchmark datasets for automated assessment.

Additionally, systematically evaluating the efficiency of human-AI collaboration requires conducting user studies involving skilled experts, such as experienced biologists and biocurators, or semi-experts, such as graduate students in biology-related fields with appropriate training. To ensure a rigorous evaluation, it is crucial to test various scenarios, including annotations performed exclusively by AI, those completed solely through manual human curation, and those achieved through human-AI collaboration.

Despite its limitations, GeneWhisperer serves as a pioneering study that leverages AI agents and advanced AI methods to assist humans in curating gene function annotations from diverse sources. Future improvements to GeneWhisperer could incorporate ongoing advancements in AI, such as LLM agents with enhanced tool-usage capabilities (Shi et al. 2024; Yuan et al. 2024). Additionally, integrating multiple agents into a cohesive multi-agent system could further enhance its utility. For example, one agent, like GeneWhisperer, could focus on curating information from various resources, another agent could synthesize this information into a formatted report, and a third agent could educate users on effectively interacting with the system.

## 6 Conclusions

To summarize, GeneWhisperer offers a user-friendly conversational system tailored for biological researchers and curators, addressing the pressing need for software support in analyzing data from multiple sources during the manual curation process. In future work, we will continue to refine GeneWhisperer to enhance its effectiveness, user interface, and broaden its scope to encompass a wider range of manual curation tools. We are also collaborating with psychologists on user studies to better understand human-AI collaboration and the human factors involved, which will be instrumental in further developing GeneWhisperer.

## Supplementary information

Supplementary materials supporting this study are available online alongside the published article.

## Declarations

### Funding

No funding was received for conducting this study.

### Competing Interests

The authors declare that they have no conflict of interest.

### Ethics approval and consent to participate

Not applicable.

### Consent for publication

Not applicable.

### Data and code availability

All data supporting the findings of this study are included in the manuscript and the supplementary materials. The study makes use of the OpenAI API with customized prompts. All prompts and relevant implementation details are provided in the manuscript and supplementary materials.

### Author contribution

XL, AW, MM and RS conceived and designed the study. XL and MF conducted the experiments. All authors (XL,AW, MM, SCA, XD, MF, CP, RS) contributed to data analysis, interpretation, and discussion of results. XL drafted the manuscript with input from the other authors. All authors reviewed and revised the manuscript.

## Footnotes

1 https://blast.ncbi.nlm.nih.gov/Blast.cgi

2 https://www.ucl.ac.uk/cardiovascular/research/pre-clinical-and-fundamental-science/functional-gene-annotation

## Notes

### Competing Interest Statement

The authors have declared no competing interest.

### Summary of Updates

This version of the manuscript includes no changes to the scientific content. The only modification is a change in formatting: we have updated the document from a two-column layout to a single-column format.

