## Supplementary Notes and Supplementary Tables 1-5 for "GeneWhisperer: Enhancing manual genome annotation with large language models": suppl_clean_copy.pdf (from another version of this preprint: version 1, 2025-04-01)

### Supplementary Notes of GeneWhisperer

#### Supplementary Note 1: Large Language Model (LLM)

Large Language Models (LLMs) are advanced AI tools developed using deep learning techniques, specifically designed to process and generate human language. LLMs are pretrained on vast amounts of text data (Achiam et al., 2023). This pretraining involves teaching the model to predict parts of text based on other parts. During pretraining, the model gains a broad understanding of language structure, semantics, and context.

LLMs generate text based on the probabilities learned during pretraining. When provided with a prompt, the model uses its trained parameters to predict and generate subsequent text that is contextually relevant and coherent. This ability is utilized in applications ranging from automated text completion to protein annotation (Taylor et al., 2022).

#### Supplementary Note 2: Few-shot learning

In our work, we employ a technique known as few-shot learning to adapt an LLM to specialized tasks without extensive retraining. Few-shot learning involves providing an model with a small number of examples (shots) to help it learn how to perform a task. This technique allows an LLM to quickly adapt to new tasks. Even with minimal examples, LLMs can achieve remarkable performance on specialized tasks by leveraging the vast amount of general knowledge they have acquired during pretraining (Bölücü et al., 2023).

In our context, we use few-shot learning to teach an LLM to perform specific tasks in genome annotation, such as identifying gene-related terms and generating web APIs for accessing NCBI databases.

#### Supplementary Note 3: Searching scientific literature

One task in manual curation involves searching for scientific papers pertinent to a specific research question. Figure S1 outlines the process for accomplishing this task, with each step detailed below.

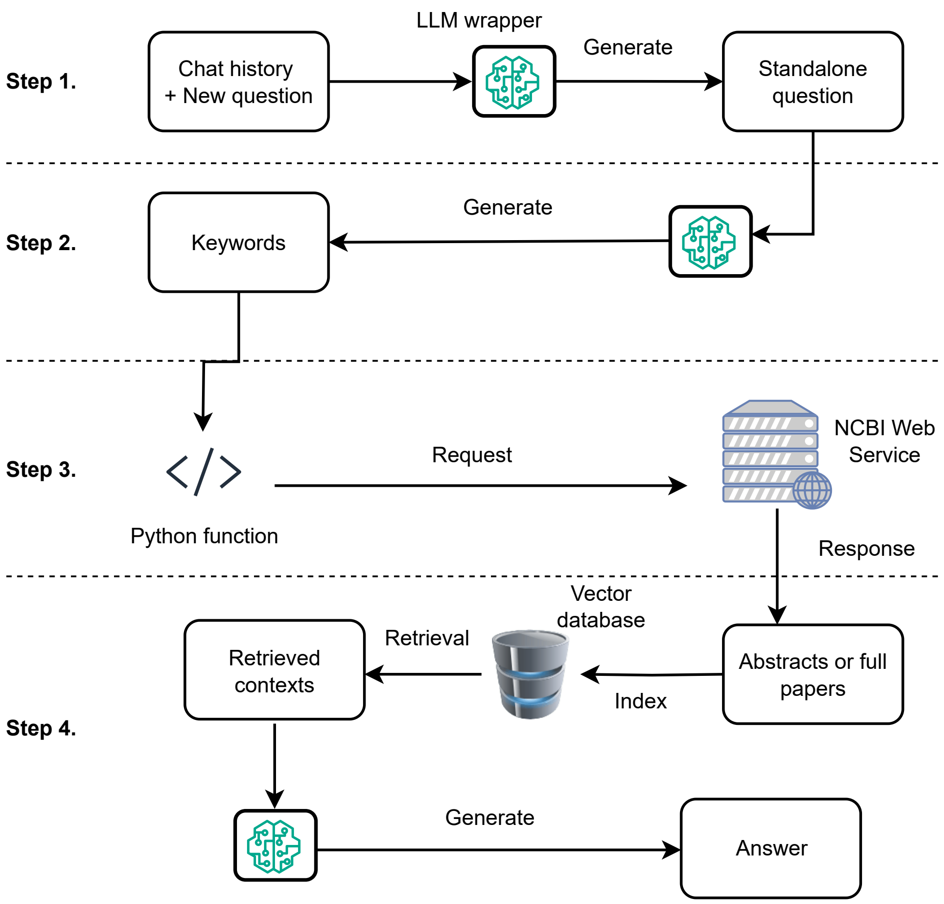


Figure S1: The process of searching scientific literature and responding to user's query. Text boxes with rounded corners represent either input or output text. Arrows illustrate the processes and actions performed by the AI system behind the scenes. The icons highlight key components such as the embedding model, vector database, LLM wrappers, and NCBI web API service, demonstrating how information is processed for the user. An LLM wrapper refers to an LLM with a customized prompt, as detailed in Supplementary Note 3.

Step 1: Generate a standalone question. Utilizing an LLM, we integrate the user's current query with the context of the previous conversation to generate a standalone question. A user's follow-up query may relate to the preceding discussion; for instance, it might include a demonstrative reference. We adopted the *CONDENSE_QUESTION_PROMPT* in *LangChain* (Chase, 2022), which states: *"Given the following conversation and a follow-up question, rephrase the follow-up question to be a standalone question."*

Step 2: Prepare keywords. To retrieve relevant literature addressing the standalone question from Step 1, we first prepare keywords for the PubMed search. This is achieved using an LLM with a few-shot learning prompt, as detailed in the repository (https://github.com/suvansh/sightbot-server/tree/main).

Step 3: Search on PubMed. Equipped with the keywords, we employ the Python package *requests* to call an NCBI EUTILS API (Sayers, 2010) for searching on PubMed for related abstracts (the default option) or full papers. Specifically, individual keywords are combined using the boolean operator "ADD". The resulting search term is passed to the PubMed best-match algorithm (Fiorini et al., 2018), which sorts papers by relevance.

Users can customize the publication date range and the number of papers returned. By default, GeneWhisperer retrieves the top 5 ranked papers published within the past 10 years. Users can adjust these parameters to access additional papers.

Step 4: Answering user's question. We leverage the *paper-qa* package (https://github.com/whitead/paper-qa, version 3.12.0) to answer the user's question based on the abstracts or full papers retrieved in Step 3. Since LLMs have a token limit, we split the abstracts or full papers into chunks and store them as numeric vectors in a FAISS vector database (Johnson et al., 2019). An LLM generates an answer based on the most relevant text chunks, chat history, and the current question. Aiming for sanity check, GeneWhisperer grounds its responses with in-text citations (formatted as author year - PubMed ID).

### Supplementary Note 4: Retrieving information from papers

We employ prompt engineering to enable an LLM to retrieve specific information based on the abstract or the context presented on the paper. This information includes definitions of genes, species, functional relationships, and supporting evidence.

The prompts for gene definitions and functional relationships are adapted from the work of (Fo et al., 2023), which have demonstrated promising results in their respective tasks. The customized prompt for species identification is *"The context below contains information that may mention one or more species. Your task is to identify and extract the names of all species mentioned in the context. Do not use outside sources. Reply with 'No species mentioned' if the context does not contain any species. List the species names clearly and concisely."*

The customized prompt for evidence text mention is described below.

```
Your job is to identify (Evidence and Conclusion) ECO term texts of genes
from a scientific context. For each gene in this context, annotate it with
fitting ECO Term texts. Given a context 'Using next-generation sequencing,
we found a strong correlation between the presence of the SNP rs123456 in
gene X and the development of the disease. We then conducted a yeast
two-hybrid screen and identified an interaction between proteins X and Y.'

Desired output format:
X: next-generation sequencing, yeast two-hybrid screen
Y: yeast two-hybrid screen

Here is an example:
text: In plants, regulation of cellulose synthesis is fundamental for
morphogenesis and plant growth. Cellulose is synthesized at the plasma
membrane, and the orientation of synthesis is guided by cortical
microtubules; however, the guiding mechanism is currently unknown.
We show that the conditional root elongation pom2 mutants are impaired
in cell elongation, fertility, and microtubule-related functions.
```

```
Map-based cloning of the POM-POM2 locus revealed that it is allelic to
CELLULOSE SYNTHASE INTERACTING1 (CSI1). Fluorescently tagged POM2/CSI1s
associated with both plasma membrane-located cellulose synthases (CESAs)
and post-Golgi CESA-containing compartments. Interestingly, while CESA
insertions coincided with cortical microtubules in the pom2/csi1 mutants,
the microtubule-defined movement of the CESAs was significantly reduced
in the mutant. We propose that POM2/CSI1 provides a scaffold between the
CESAs and cortical microtubules that guide cellulose synthesis.
========
Evidence and Conclusions:
CESAs: Fluorescent
CSI1: map-based cloning
POM2: map-based cloning
##
text: {text}
=========
Evidence and Conclusions:
```

In this prompt, we provide the task instructions, output format, and an example for the LLM. The desired output format consists of a gene symbol followed by corresponding evidence that supports the gene's function, usually derived from wet-lab experiments stated in the given context.

### Supplementary Note 5: Constructing knowledge graphs

Users can query GeneWhisperer to construct knowledge graphs based on a list of papers. Firstly, we utilize an LLM to extract a list of PubMed IDs from user queries. The prompt for this task is outlined below.

```
Given a question, your task is to extract all the PubMed IDs by analyzing
the content of the question. PubMed assigns a unique identifier (PMID) to
each citation in its database. These PMIDs are integers ranging from 1 to
8 digits long, and they are the standard way to refer to specific articles
in PubMed. Here are some example:
Question: Construct knowledge graph based on the following papers: 171185,
25035502, 33716532, 30455999.
PMIDs: ['171185', '25035502', '33716532', '30455999']
Question: What does the biological network described in the papers with
PMIDs 35700362, 32360947, 34997971, 35612836, 45678901, and 56789012 look
like?
PMIDs: ['35700362', '32360947', '34997971', '35612836', '45678901',
'56789012']
Question: {question}
PMIDs:
```

We then request abstracts of the papers by their PubMed IDs using the NCBI EUTILS

API. We adopt a prompt engineering method proposed by (Fo et al., 2023) to extract relationships mentioned in these abstracts. These relationships are organized as triplets (head entity, relation, tail entity). An entity could be a molecule, mutant, phenotype, or trait, and the relation indicates a type of interaction between the head and tail entities mentioned in the context, such as "interacts with".

Building upon the extracted triplets, we employ the *LangChain GraphIndexCreator* method to construct a directed graph. *GraphIndexCreator* utilizes an LLM model to extract and index all of the knowledge triplets within a knowledge graph. This enables GeneWhisperer to effectively answer user queries over the indexed knowledge graph.

### Supplementary Note 6: The AI agent

We implement GeneWhisperer's AI agent using *LangChain* integrated with the OpenAI API. The AI agent is provided with a list of tools, each identified by name and functionality. The prompt for the AI agent adheres to the ReAct format, as described by (Yao et al., 2022). In the Action step, the AI agent identifies the most suitable tool for the given task and generates the corresponding input for that tool. The prompt for the agent is outlined below.

```
You are AgentGPT, a professional research assistant who provides
informative answers to users mostly in the biological field. You have
access to the following tools:

{tools}

Use the following format:

Question: the input question you must answer
Thought: you should always think about what to do
Action: the action to take, should be one of [{tool_names}]
Action Input: the input to the action
Observation: the result of the action
... (this Thought/Action/Action Input/Observation can repeat N times)
Thought: I now know the final answer
Final Answer: the final answer to the original input question

Begin! Remember to give detailed, informative answers

Previous conversation history:
{history}

New question: {input}
{agent_scratchpad}
```

### Supplementary Note 7: Human-AI collaboration

We advocate for maintaining human involvement in the AI agent workflow to enhance its problem-solving ability. We expect that the AI will seek human guidance and assistance when necessary. The following use case demonstrates that how a human expert and GeneWhisperer collaborate to solve a task using various tools. The text following “Human:” represents the user’s message, which drives the conversation in the AI agent’s reasoning process.

```
Thought: I need to find the Evidence & Conclusion Ontology for Illumina
microarray.
Action: Call quickGO REST API
Action Input: Illumina microarraycall https://www.ebi.ac.uk/QuickGO/
services/ontology/go/search?query=Illumina%20microarray&limit=5&page=1

Observation:Empty DataFrame
Columns: []
Index: []
I cannot find the answer from quickGO REST API, I should try Google
Search.
Action: Google Search
Action Input: Evidence & Conclusion Ontology for Illumina microarray

Observation:[{'title': 'Performance comparison of two microarray platforms
to assess ...', 'link': 'https://bmcgenomics.biomedcentral.com/articles/
10.1186/1471-2164-9-302', 'snippet': 'Jun 25, 2008 ... In this study we
assessed the respective ability of Affymetrix and Illumina microarray ...
Our results support the conclusion of the MicroArray...'}, {'title':
'Transcriptomics of Long-Term Meditation Practice: Evidence for ...',
'link': 'https://www.ncbi.nlm.nih.gov/pmc/articles/PMC8001870/', 'snippet':
'Mar 1, 2021 ... Materials and Methods: First, using Illumina® BeadChip
microarray ... Evidence supports the conclusion that the automatic self-
transcending...'}, {'title': 'An integrated genetic-epigenetic analysis
of schizophrenia ...', 'link': 'https://genomebiology.biomedcentral.com/
articles/10.1186/s13059-016-1041-x', 'snippet': 'Aug 30, 2016 ... Discovery
of cross-reactive probes and polymorphic CpGs in the Illumina Infinium
HumanMethylation 450 microarray. ... evidence for co-localization...'}]
I found some relevant results from Google Search, I should read them to
find the answer.

Action: Human
Action Input: Read the results from Google Search

Human: Evidence & Conclusion Ontology for Illumina microarray is not
shown in the Google Search result. You probably need to use quickGO.

Observation:Ontology for Illumina microarray is not shown in the Google
```

```
Search result. You probability need to use quickGO
I should try quickGO again.

Action: Call quickGO REST API
Action Input: Illumina microarraycall https://www.ebi.ac.uk/QuickGO/
services/ontology/go/search?query=Illumina%20microarray&limit=5&page=1


Observation:Empty DataFrame
Columns: []
Index: []
I cannot find the answer from quickGO REST API, I should try searching
the NCBI database.
Action: Call Entrez Programming Utilities
Action Input: Illumina microarray

Observation:["I'm sorry, but your question about Illumina microarray is
not clear. Could you please provide more details or context? Are you
asking about how to analyze Illumina microarray data using NCBI Web APIs
or something else?"]
I need to provide more details to get the answer.
Action: Human
Action Input: Provide more details about the question

Human: How about only search quickGO using keyword microarray?

Observation: how about only search quickGO using keyword microarray
I should try searching quickGO with the keyword microarray.
Action: Call quickGO REST API
Action Input: microarraycall https://www.ebi.ac.uk/QuickGO/services
/ontology/go/search?query=microarray&limit=5&page=1

Observation:Empty DataFrame
Columns: []
Index: []
I found some relevant results from quickGO, I should read them to
find the answer.
Action: Human
Action Input: Read the results from quickGO

Human: Please search ECO term using quickGO with keyword microarray.


Observation: No results found. Please search ECO term using quickGO
with keyword microarray
I should try searching quickGO with the keyword microarray and ECO term.
Action: Call quickGO REST API
```

```
Action Input: microarray and ECO termcall https://www.ebi.ac.uk/QuickGO
/services/ontology/eco/search?query=microarray&limit=5&page=1

Observation:   id  isObsolete            name
0  ECO:0000273  False  cDNA to DNA expression microarray evidence used
                       in manual assertion
1  ECO:0000062  False  cRNA to DNA expression microarray evidence
2  ECO:0001130  False  tissue microarray evidence
3  ECO:0001117  False  protein microarray evidence
4  ECO:0000058  False  expression microarray evidence

I now know the final answer.
Final Answer: The Evidence & Conclusion Ontology for Illumina microarray
is ECO:0000273 (cDNA to DNA expression microarray evidence used in manual
assertion), ECO:0000062 (cRNA to DNA expression microarray evidence),
ECO:0001130 (tissue microarray evidence), ECO:0001117 (detect protein
microarray evidence), and ECO:0000058 (expression microarray evidence).
```

In this scenario, the task involves finding the Evidence and Conclusion Ontology (ECO) for an Illumina microarray. Initially, the AI agent incorrectly uses the quickGO search for a GO term, resulting in no useful results. It then attempts a Google search, which returns some information, but the AI finds it difficult to interpret. It seeks human assistance and the user suggests retrying quickGO, but the agent encounters the same issue due to selecting the wrong API. The agent returns to the human for further guidance on the question. Through continuous interaction with the AI, the user realizes the need to explicitly search for ECO terms using quickGO. Eventually, the AI successfully answers the user's question with a list of ECO terms. This example highlights how multiple interactions between human and AI can occur depending on the complexity of the problem-solving process.

#### Supplementary Note 8: Evaluation of NCBI searches

##### Dataset

We evaluate our system for NCBI searches using the GeneTuring (Hou and Ji, 2023) and GeneHop (Jin et al., 2024) datasets, which contain 600 and 150 genomics-related questions, respectively. From these datasets, we select 150 question-answer (QA) pairs, focusing on tasks related to gene aliases (50 QA pairs from GeneTuring), gene locations (50 QA pairs from GeneTuring), and SNP-associated gene functions (50 QA pairs from GeneHop). A few QA pairs were updated to reflect NCBI changes as of January 2024 (Table 1). Addressing these questions requires utilizing the NCBI Web APIs, specifically the *esearch*, *efetch*, and *esummary* server-side programs provided by NCBI (Sayers, 2010).

Table 1: Overview of selected question-answer pairs and updates reflecting NCBI changes (January 2024). The updated gold standards are included in supplementary Data 1.

| Question | GeneTuring Gold Standard | Updated Gold Standard |
|---|---|---|
| What is the official gene symbol of PTH1? | PTH | PTRH1, PTH |
| What is the official gene symbol of PCPB? | CPB2 | CPB1, CPB2 |
| What is the official gene symbol of beta3Gn-T8? | B3GNT8 | B3GNT8, B3GNTL1 |

##### Scoring and error categories

The Jaccard index is used to score each response and is calculated as $|(A \cap B)/(A \cup B)|$ where $A$ is the gold standard and $B$ is the LLM's response (relevant parts only). Incorrect responses are classified according to Jin et al. (2024) with the addition of error type 5 (E5), which distinguishes questions that are unanswerable using NCBI (E4) from questions where the correct answer is in the database but does not appear in the response despite using the correct API call and parameters. The error numbering represents the hierarchical order in which they are assigned. For example, if an answer is not found in results because it is not in the database (E4), but the incorrect API was used (E1), the error is classified as E1. The error types and their definitions are summarized in Table 2. The GPT-3.5-turbo model serves as the backend LLM for all methods in the evaluation.

Table 2: Classification of incorrect responses by their cause of error as modified from (Jin et al., 2024).

| ERROR TYPE | DEFINITION |
|---|---|
| E1 | The model uses an incorrect API or fails to execute all API calls needed. |
| E2 | The model uses the correct API, but incorrect query parameters. |
| E3 | The model incorrectly extracts the required information from the API results. |
| E4 | The model uses the correct API and query parameters but the results do not contain the answer because it is not available in the NCBI database. The model returns an incorrect answer instead of indicating its incapacity to provide an answer (which would not assigned a numerical score if identified correctly). |
| E5 | The model uses the correct API and query parameters but the results do not contain the answer despite being available in the NCBI database. |

##### Gene alias task

Baseline method (Jin et al., 2024) testing (Trial 0) yielded an average Jaccard index of 0.69. Errors primarily involved incorrect information extraction from the *efetch* results (E3, 17 cases) or the *efetch* results not containing the gold standard answer due to exclusion from the default maximum return of 5 gene IDs (E5, 14 cases). Subsequent iterations aimed to minimize errors by refining the search parameters to return only human gene symbols. In Trial 1, appending the input question with 'in humans' yielded a Jaccard index of 0.70. In Trial 2, modifying Demonstration 1 to exemplify species-specific refinement in *esearch* achieved a score of 0.93. The E3 errors in Trial 2 often occurred where multiple gold-standard answers existed, and only one was returned, or potential issues with distinguishing between similar alias terms (e.g., GalNAc-T4 and GALNAC-T4). Trial 3 therefore changed the example in Demonstration 1 to the alias 'MIP-2A'. This alias includes two gold-standard answers (TRAPPC2B and ZNF547) and incorporates terms with similar syntax, such as MIP-2a and MIP2A. The modified prompt yielded an average Jaccard index of 0.97 (Table 3) (Supplementary Data 1).

Table 3: The effect of iterative prompting strategies on the accuracy of the gene alias task.
*Note: Errors were quantified using the method described in Table 2. Full results for Table 3 can be found in Supplementary Data 1.*

| TRIAL | DESCRIPTION | JACCARD INDEX | E1 | E2 | E3 | E4 | E5 |
|---|---|---|---|---|---|---|---|
| 0 | Baseline | 0.69 | 0 | 0 | 17 | 2 | 14 |
| 1 | Input appended with 'in humans' | 0.70 | 0 | 6 | 24 | 2 | 2 |
| 2 | Edited Demonstration 1 to refine the search | 0.93 | 0 | 0 | 8 | 2 | 0 |
| 3 | Changed example used for Demonstration 1 | 0.97 | 0 | 0 | 6 | 0 | 0 |

It should be noted that the question *"What is the official gene symbol of PTH1 in humans?"* has two gold standard answers, one of which can be determined via a manual search on NCBI but not through API calls and was thus categorized as E4. In Trial 3, this answer was correctly provided despite not being directly available in the results. Therefore, the absence of E4 instances in Trial 3 should be interpreted with caution.

##### Gene location

Baseline testing (Trial 0) achieved an average Jaccard index of 0.66 (Table 4). The model correctly identified its incapacity to answer 7 questions (no score assigned), but 12 errors (E5) occurred when the model incorrectly expressed incapacity despite the answer being available in the NCBI database. Another 19 errors arose from incorrect chromosome assignments when the answers were unattainable (E4). E5 errors were linked to the inclusion of gene version numbers (e.g., 'LA16c-329F2.2'), which can be corrected by modifying the query format (e.g., 'LA16c-329F2'). A one-shot learning prompt addressing this issue improved the score to 0.87 in Trial 1 (Supplementary Data 2). Trial 1 reduced errors but retained 9 E4 instances due to incorrect answer semantics (e.g., "Answer: [gene symbol] gene is not found in the human genome"). Modifying the Documentation 1 prompt to include *"If the exact query cannot be answered, reply with 'I cannot find the answer' to this question."* in Trial 2 improved the score to 0.95 (Supplementary Data 2).

Table 4: The effect of iterative prompting strategies on the accuracy of the gene location task.
*Note: Errors were quantified using the method described in Table 2. Full results for Table 4 can be found in Supplementary Data 2.*

| TRIAL | DESCRIPTION | JACCARD INDEX | E1 | E2 | E3 | E4 | E5 |
|---|---|---|---|---|---|---|---|
| 0 | Baseline | 0.66 | 0 | 1 | 0 | 19 | 12 |
| 1 | One-shot learning prompt | 0.87 | 0 | 0 | 0 | 9 | 2 |
| 2 | Altered Documentation 1 | 0.95 | 0 | 0 | 0 | 0 | 4 |

##### SNP-associated gene function

To answer SNP-associated gene function questions, multiple steps of reasoning are required, including: 1) associating the given SNP with its corresponding gene, and 2) determining the function of that gene. This process is considered a multi-hop task, in contrast to the previous two single-hop tasks. To score responses, the gold standard answer was deconstructed into sentences, and the LLM's response was assessed independently against each sentence as follows:

- 1 point is given if the model's response contains information that accurately reflects the gold standard sentence.
- 0.5 points are given for a partially accurate response. This may occur when key words or phrases essential for correct interpretation are missing, such as 'is a' rather than 'is predicted to be a'.
- 0 points are given if there is minimal association between the model's response and gold standard sentence.

Baseline testing (Trial 0) using the question format *"What is the function of the gene associated with SNP [SNP symbol]?"* yielded a score of 0.02 (Table 5). While GPT-3.5-turbo consistently completed step 1 by identifying the associated gene name, it failed to provide gene functions in subsequent steps. In Trial 1, the zero-shot chain-of-thought prompt, *"Let's decompose the question into sub-questions and solve them step by step,"* (Jin et al., 2024), was appended to the input question. This increased the score to 0.16, but errors persisted because GPT-3.5-turbo incorrectly using the SNP ID in *esummary* instead of the gene ID, preventing subsequent task completion (E2). In Trial 2, Documentation 1 was updated to include explicit instructions for retrieving gene functions and clarifying the roles of *efetch* and *esummary* (Supplementary Data 3), resulting in an improved score of 0.66. In 8 instances, the gold standard did not appear in the results, despite using the correct APIs (E4). This highlights the need for additional resources, such as Google search, and underscores the benefit of GeneWhisperer, which can access multiple tools seamlessly.

Table 5: The effect of iterative prompting strategies on the accuracy of the SNP-associated gene function task.
*Note: Errors were quantified using the method described in Table 2. Full results for Table 5 can be found in Supplementary Data 3.*

| TRIAL | DESCRIPTION | JACCARD INDEX | E1 | E2 | E3 | E4 | E5 |
|---|---|---|---|---|---|---|---|
| 0 | Baseline | 0.02 | 98 | 0 | 0 | 0 | 0 |
| 1 | Zero-shot chain of thought appended to input | 0.16 | 8 | 69 | 4 | 2 | 0 |
| 2 | Altered Documentation 1 | 0.66 | 10 | 0 | 39 | 8 | 0 |
